# An Automated System for Examining Target Chasing and Interception in Rodents

**DOI:** 10.64898/2026.11.24.769042

**Authors:** Xiaoxiao Lin, Jarod Bussey, Chancellor M. Crawford, Miranda De Leon, Grace Fisher, Luehan Wang, Jingtang Sun, Andrew S. Alexander

## Abstract

Rodents exhibit sophisticated navigation behaviors in both naturalistic and laboratory settings, including the pursuit and interception of moving targets during predation, play, and social interactions. Prior work has shown that rodents engage in pursuit of purely visual moving targets for reward. However, limited experimental control over target movement in existing approaches makes it difficult to systematically examine how animals adapt their pursuit strategies to changing target dynamics. To address this challenge, we developed an automated behavioral platform for studying visually guided pursuit and interception in rodents under precisely controlled conditions. Specifically, we introduce a behavioral-closed-loop physical system that combines live animal tracking with programmable visual targets presented via floor projection, and automated trial control, reducing manual intervention while separating pursuit into distinct behavioral phases for quantitative analysis. A custom Python-based GUI allows users to generate and modify target trajectories, while a Bonsai workflow brings experimental control and synchronization into a single software environment. We provide detailed instructions for constructing the system, configuring the software, and training animals to pursue and intercept visual targets. Together, these procedures provide a reproducible and adaptable approach for investigating behavioral strategies underlying visually guided pursuit behavior.

## INTRODUCTION

Spatial navigation is a central topic in systems neuroscience, yet common rodent navigation paradigms (e.g., free foraging or track running) typically take place under relatively static conditions that lack the complex dynamics present in the natural world. The limited ethological validity of these reduced experimental conditions restrict study of the full repertoire of complex navigation behaviors that arise in real-world environments.

Recent work has sought to make navigation studies more naturalistic in various ways, including by introducing interactions with other moving agents such as conspecifics, predators, or prey^1–4^. Emergent chasing behaviors occur in both social and predatory contexts in rodents and have been increasingly studied in the laboratory^5–14^. Chasing involves pursuit of a moving goal and is behaviorally demanding: the animal must integrate sensorimotor and spatial information to guide continuous decisions that require coordinated motor output^15^. Hunting and social pursuit have therefore been adopted as neuroethologically inspired frameworks for examining a range of higher-order cognitive processes, including decision making, visual processing, social and emotional behavior, and spatial cognition. These studies have yielded important insights into the neural circuitry and computations underlying chase behavior, but have typically traded control over target movement for ease of implementation. Precision in target movement is critical for developing formalized models of animal behavior under pursuit conditions. This is especially critical for investigation of neural computations underlying emergent predictive strategies that occur during pursuit.

Here, we present a novel experimental system for studying visually-guided chasing behaviors. We harness the inclination of animals to hunt stimuli through the use of visual signals^2^. Heavily inspired by prior instantiations of a floor projection maze^16^, we use rear projection onto a behavioral arena to display a dot that follows consistently rendered, pre-generated trajectories that the mice can intercept for reward. To help others initiate their own experiments, we developed trajectory generation software which constructs ‘random’ target paths by sampling from underlying angular and linear probability distributions and concatenating the subsequent vectors. In this protocol we provide a technical description of an automated visual pursuit behavioral rig so that other laboratories may reproduce this ecosystem for their own studies (**Table 1**).

**Table 1.**
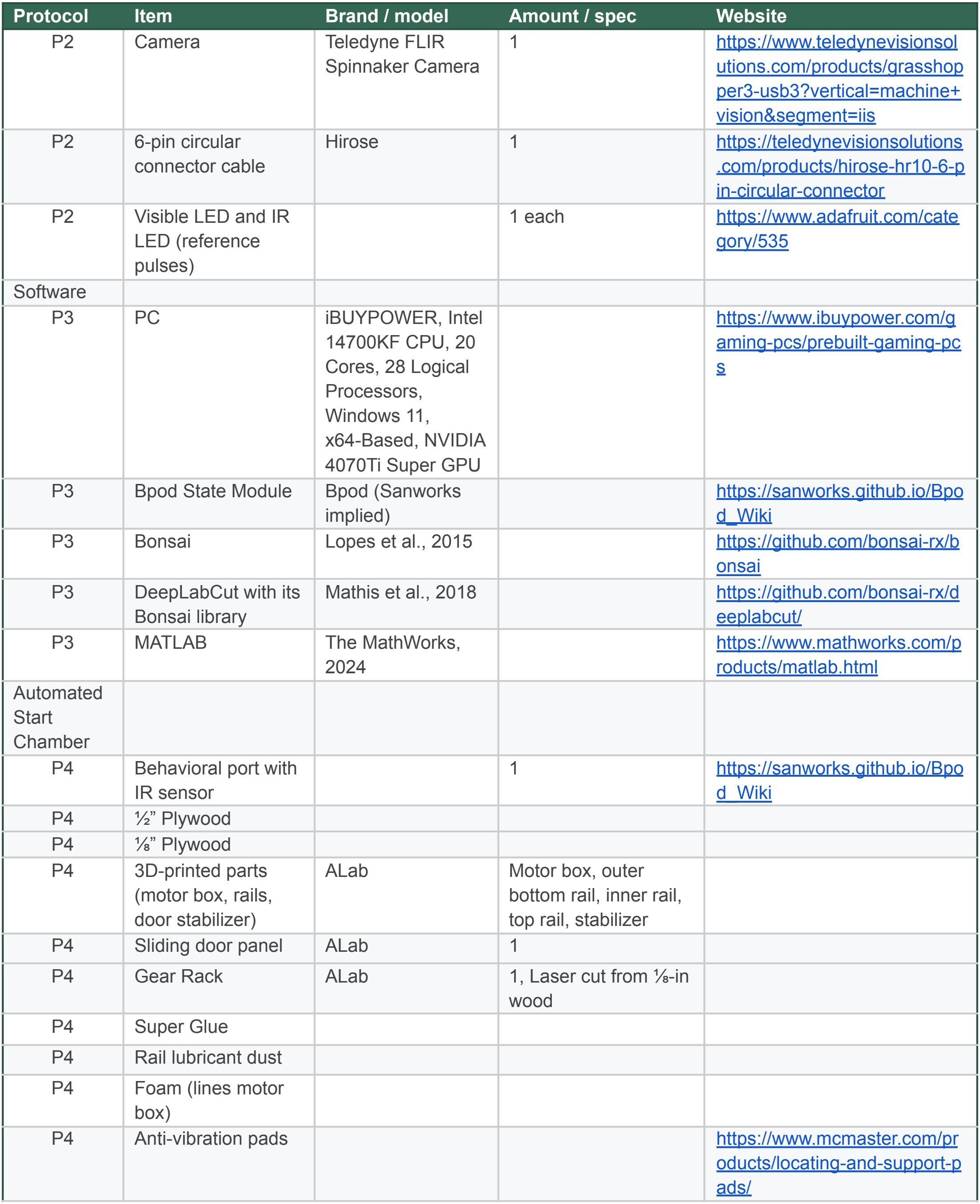
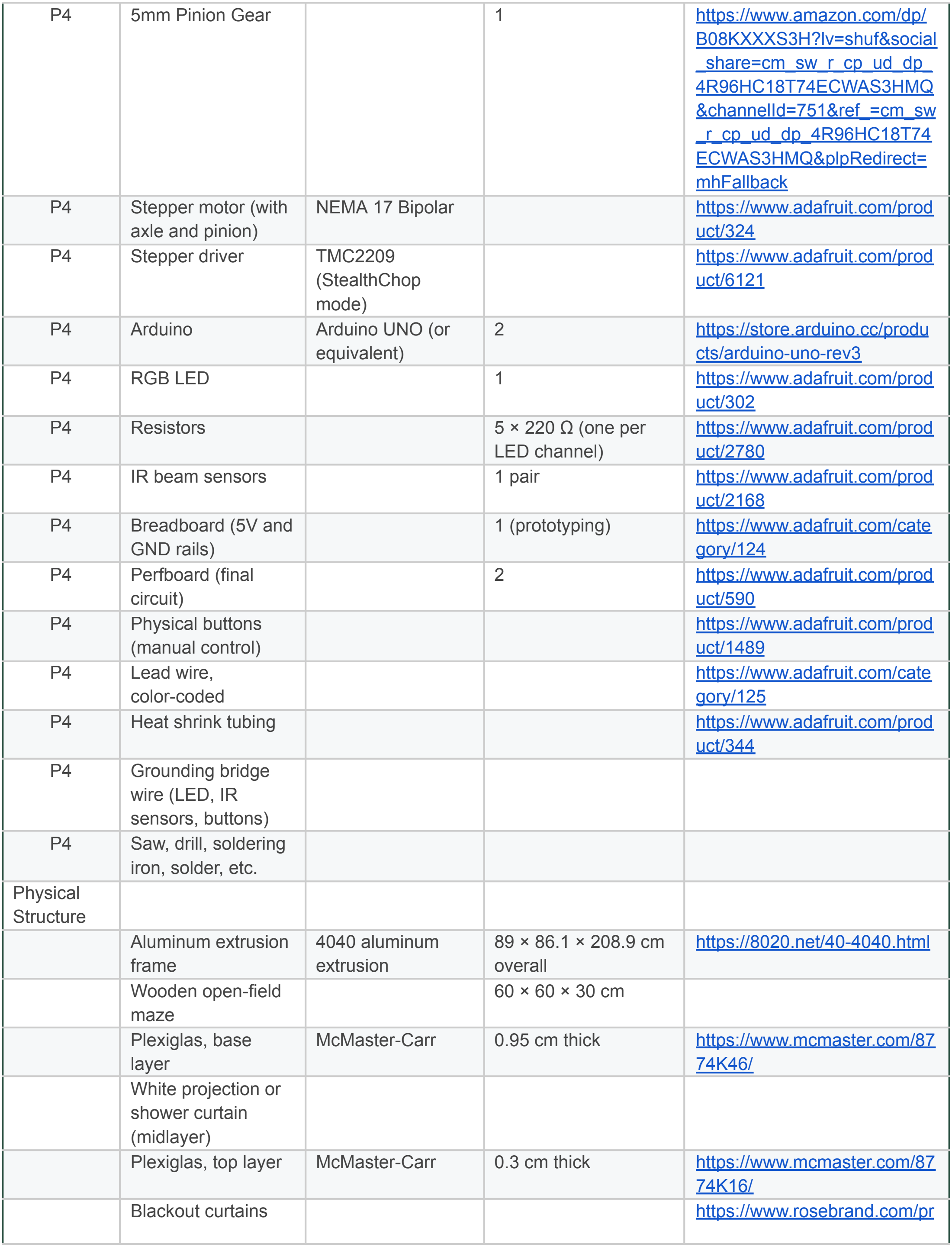
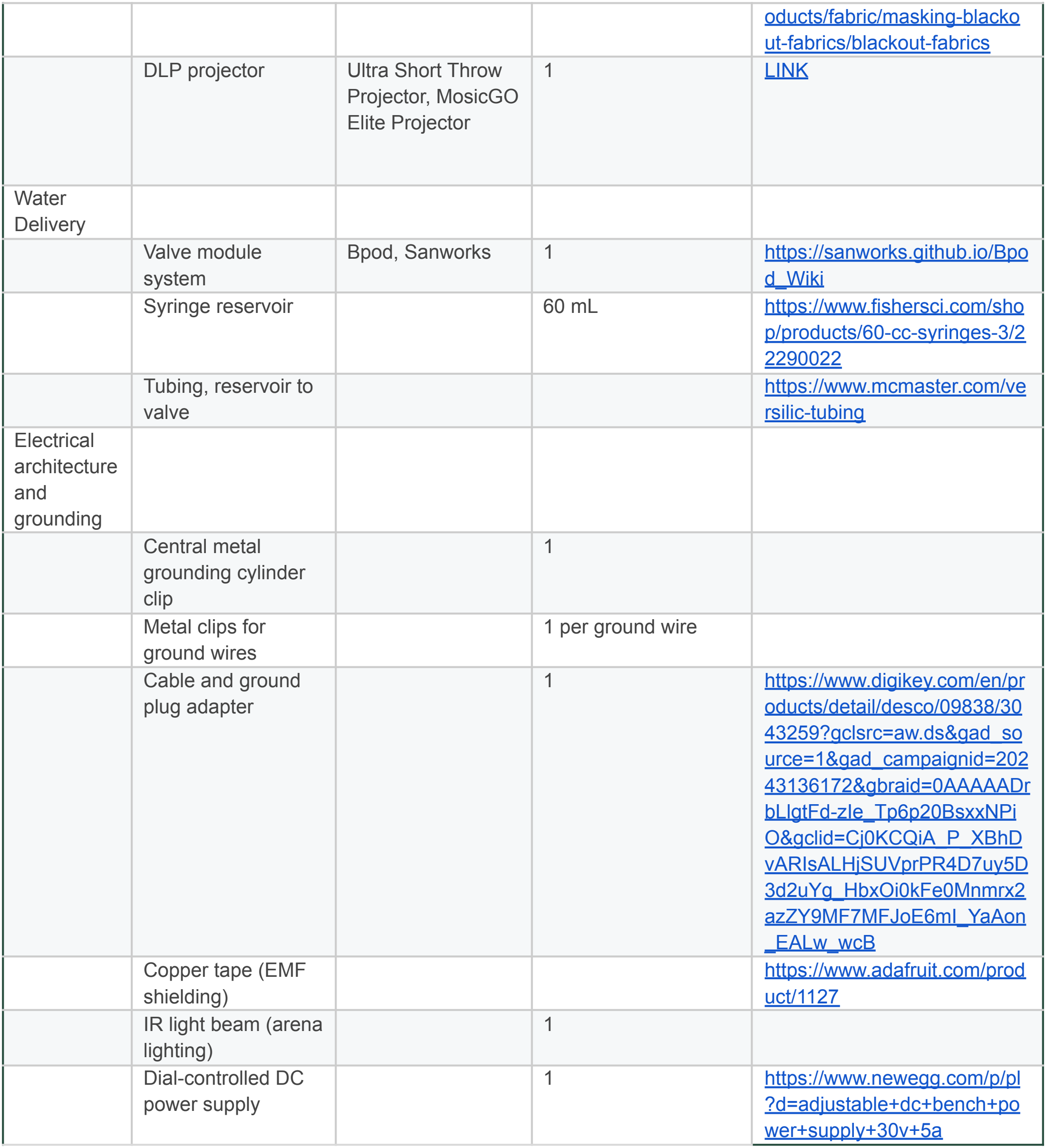
Materials List.

| Protocol | Item | Brand / model | Amount / spec | Website |
|---|---|---|---|---|
| P2 | Camera | Teledyne FLIR Spinnaker Camera | 1 | https://www.teledynevisionsolutions.com/products/grasshopper3-usb3?vertical=machine+vision&segment=iis |
| P2 | 6-pin circular connector cable | Hirose | 1 | https://teledynevisionsolutions.com/products/hirose-hr10-6-pin-circular-connector |
| P2 | Visible LED and IR LED (reference pulses) |  | 1 each | https://www.adafruit.com/category/535 |
| Software |  |  |  |  |
| P3 | PC | iBUYPOWER, Intel 14700KF CPU, 20 Cores, 28 Logical Processors, Windows 11, x64-Based, NVIDIA 4070Ti Super GPU |  | https://www.ibuypower.com/gaming-pcs/prebuilt-gaming-pcs |
| P3 | Bpod State Module | Bpod (Sanworks implied) |  | https://sanworks.github.io/Bpod_Wiki |
| P3 | Bonsai | Lopes et al., 2015 |  | https://github.com/bonsai-rx/bonsai |
| P3 | DeepLabCut with its Bonsai library | Mathis et al., 2018 |  | https://github.com/bonsai-rx/deeplabcut/ |
| P3 | MATLAB | The MathWorks, 2024 |  | https://www.mathworks.com/products/matlab.html |
| Automated Start Chamber |  |  |  |  |
| P4 | Behavioral port with IR sensor |  | 1 | https://sanworks.github.io/Bpod_Wiki |
| P4 | ½” Plywood |  |  |  |
| P4 | ⅛” Plywood |  |  |  |
| P4 | 3D-printed parts (motor box, rails, door stabilizer) | ALab | Motor box, outer bottom rail, inner rail, top rail, stabilizer |  |
| P4 | Sliding door panel | ALab | 1 |  |
| P4 | Gear Rack | ALab | 1, Laser cut from ⅛-in wood |  |
| P4 | Super Glue |  |  |  |
| P4 | Rail lubricant dust |  |  |  |
| P4 | Foam (lines motor box) |  |  |  |
| P4 | Anti-vibration pads |  |  | https://www.mcmaster.com/products/locating-and-support-pads/ |
| P4 | 5mm Pinion Gear |  | 1 | https://www.amazon.com/dp/B08KXXXS3H?lv=shuf&social_share=cm_sw_r_cp_ud_dp_4R96HC18T74ECWAS3HMQ&channelId=751&ref_=cm_sw_r_cp_ud_dp_4R96HC18T74ECWAS3HMQ&plpRedirect=mhFallback |
| P4 | Stepper motor (with axle and pinion) | NEMA 17 Bipolar |  | https://www.adafruit.com/product/324 |
| P4 | Stepper driver | TMC2209 (StealthChop mode) |  | https://www.adafruit.com/product/6121 |
| P4 | Arduino | Arduino UNO (or equivalent) | 2 | https://store.arduino.cc/products/arduino-uno-rev3 |
| P4 | RGB LED |  | 1 | https://www.adafruit.com/product/302 |
| P4 | Resistors |  | 5 × 220 Ω (one per LED channel) | https://www.adafruit.com/product/2780 |
| P4 | IR beam sensors |  | 1 pair | https://www.adafruit.com/product/2168 |
| P4 | Breadboard (5V and GND rails) |  | 1 (prototyping) | https://www.adafruit.com/category/124 |
| P4 | Perfboard (final circuit) |  | 2 | https://www.adafruit.com/product/590 |
| P4 | Physical buttons (manual control) |  |  | https://www.adafruit.com/product/1489 |
| P4 | Lead wire, color-coded |  |  | https://www.adafruit.com/category/125 |
| P4 | Heat shrink tubing |  |  | https://www.adafruit.com/product/344 |
| P4 | Grounding bridge wire (LED, IR sensors, buttons) |  |  |  |
| P4 | Saw, drill, soldering iron, solder, etc. |  |  |  |
| Physical Structure |  |  |  |  |
|  | Aluminum extrusion frame | 4040 aluminum extrusion | 89 × 86.1 × 208.9 cm overall | https://8020.net/40-4040.html |
|  | Wooden open-field maze |  | 60 × 60 × 30 cm |  |
|  | Plexiglas, base layer | McMaster-Carr | 0.95 cm thick | https://www.mcmaster.com/8774K46/ |
|  | White projection or shower curtain (midlayer) |  |  |  |
|  | Plexiglas, top layer | McMaster-Carr | 0.3 cm thick | https://www.mcmaster.com/8774K16/ |
|  | Blackout curtains |  |  | https://www.rosebrand.com/products/fabric/masking-blackout-fabrics/blackout-fabrics |
|  | DLP projector | Ultra Short Throw Projector, MosicGO Elite Projector | 1 | LINK |
| Water Delivery |  |  |  |  |
|  | Valve module system | Bpod, Sanworks | 1 | https://sanworks.github.io/Bpod_Wiki |
|  | Syringe reservoir |  | 60 mL | https://www.fishersci.com/shop/products/60-cc-syringes-3/22290022 |
|  | Tubing, reservoir to valve |  |  | https://www.mcmaster.com/versilic-tubing |
| Electrical architecture and grounding |  |  |  |  |
|  | Central metal grounding cylinder clip |  | 1 |  |
|  | Metal clips for ground wires |  | 1 per ground wire |  |
|  | Cable and ground plug adapter |  | 1 | https://www.digikey.com/en/products/detail/desco/09838/3043259?gclsrc=aw.ds&gad_source=1&gad_campaignid=20243136172&gbraid=0AAAADrbLlgtFd-zle_Tp6p20BsxxNPiO&gclid=Cj0KCQiA_P_XBhDvARIsALHjSUVprPR4D7uy5D3d2uYg_HbxOi0kFe0Mnmrx2azZY9MF7MFJoE6ml_YaAon_EALw_wcB |
|  | Copper tape (EMF shielding) |  |  | https://www.adafruit.com/product/1127 |
|  | IR light beam (arena lighting) |  | 1 |  |
|  | Dial-controlled DC power supply |  | 1 | https://www.newegg.com/p/pl?d=adjustable+dc+bench+power+supply+30v+5a |

### A COMPLETE ECOSYSTEM FOR CLOSED-LOOP TARGET CHASING

We have developed a fully automated behavioral ecosystem for studying visual target pursuit in freely moving rodents (**Figure 1, Video 1**). Our primary contributions include a software package for generating moving stimuli, a BonsaiRx based workflow for presenting moving stimuli via projection^17^, and a description of an integrated hardware system for controlling trial structure and delivering reward in an automated fashion. We begin by describing the trajectory generation software and Bonsai workflow before moving to the physical apparatus, including an automated start box, and related protocols. When assembled as described here, mice can perform hundreds of pursuit trials per day with minimal supervision from the experimenter.

**Figure 1.**
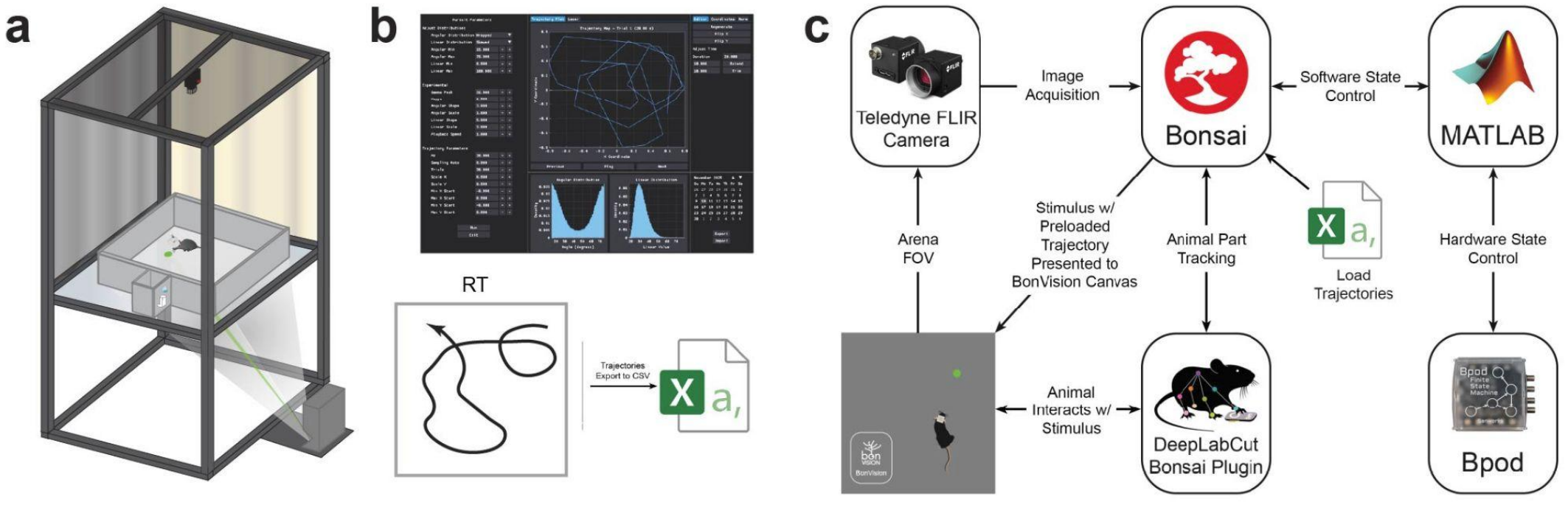
Schematic of the projection-based pursuit rig and integrated behavioral-neural recording workflow. **a.** Depiction of floor-projection system displaying a simulation-generated moving target while a mouse navigates the arena, **b.** Trajectories are created with a custom GUI with parameters controlling angular distribution, curvature, and speed, **c.** Flow chart summarizing the full software and hardware pipeline for automated target chasing behavioral studies.

### TARGET TRAJECTORY GENERATION SOFTWARE

We have built Python software that allows for the generation of randomized dot trajectories based on predetermined angular and linear velocity distributions. Using these trajectories, a moving circular dot travels around the arena floor that animals can pursue and intercept for liquid reward. Interception is defined by tracking the mouse in real time and calculating the distance between the mouse’s nose and the known location of the moving target (discussed further below). Here we provide an overview on how to use our software to generate your own trajectories and facilitate target presentation through Bonsai.

Details on how to download, install and run the software is included in this GitHub (https://github.com/alab-ucsb/alab-pursuitSimulatorRP). To generate random trajectories, start in the main graphical user interface (GUI) of the Pursuit Simulator. All settings and parameters are described in **Figure 2** and **Protocol 1**. Trajectory generation requires defining “durations” for all sessions, the speed of target movement via the “Linear Range” parameter and angular distribution to control changes in movement direction. Clicking “Run Simulation” applies the settings and generates a full set of unique trials based on the predetermined parameters. Individual trials can be selected and inspected using the trajectory map in the center of GUI. Users can adjust the parameters for a selected trial and click the “Regeneration” button to generate a new trajectory without affecting the other trials. Preferences for what types of trajectories are necessary is dependent on experimentation, but generally few moderately sized loops (approximately 10 to 20 cm in diameter) and broad spatial coverage across the arena are recommended for mice. Once trajectory features are satisfactory, naming and clicking “Export CSV” will save the trajectory outputs to your chosen local file directory. This output file can be loaded into Bonsai to render the trajectory from the projector onto the maze floor (described further below).

**Figure 2.**
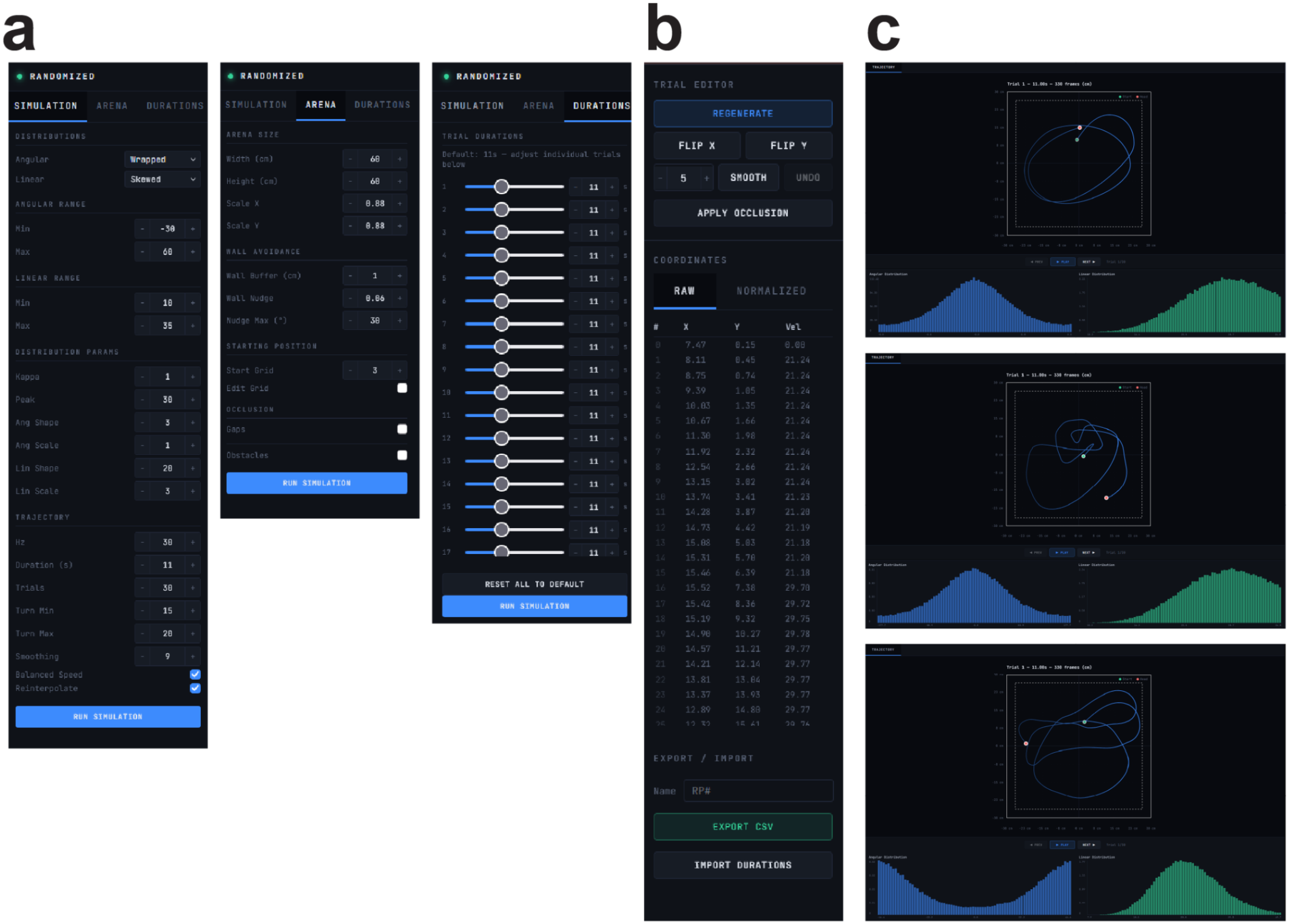
Depiction of Trajectory Generation Graphical User Interface. **a**. Input parameters for defining underlying angular and linear speed distributions that trajectories are drawn from. Details for each parameter are provided in **Protocol 1. b**. Trial editor, used for editing and regenerating individual trials after initial construction, c. Trajectory view. GUI for flipping through each trial, playing the trial as a video, and examining the speed distributions used to generate each trial (bottom). Shown are three different example trajectories drawn from speed distributions with different angular and linear speed parameters.

**Protocol 1 - Generating target trajectories in Pursuit Simulator GUI**.

*Basic operation*.

1. *Trajectory Map:*
  a. This shows the trajectory that was generated and exactly what will appear in the arena.
2. *Regenerate:*
  a. This button will regenerate the trajectory of the current trial. Keep in mind that you cannot undo a regeneration, so make sure you actually want to regenerate before doing so.
3. *Flip X and Flip Y:*
  a. Flip X will multiply all of the X-coordinates by -1, which will flip the trajectory over the y-axis.
  b. Flip Y will multiply all of the Y-coordinates by -1, which will flip the trajectory over the x-axis.
  c. You can use both Flip X and Flip Y, just click on one first and then the other.
4. *Play:*
  a. View a video of the trajectory so you can see what it will look like after each frame. You can click on ‘Laser’ at the top of the plot to visualize it as a green dot.
  b. *Playback Speed*: speed at which the trajectory appears when you press “Play” in the generator – does not affect the actual trajectory.
5. *Previous and Next:*
  a. Navigate trial order sequentially.
6. *Export:*
  a. Export the trajectory as a csv. The file will be saved in the ‘Trajectory Generators’ folder in a folder called “trajectories” as “trial_results_MM-DD-YY.csv”
  b. Important: If you have already generated a file in the same day, clicking on this again will overwrite that file.
7. *Trials:*
  a. Define the amount of trials. The default is 30 trials. The minimum amount is 1, and there is no maximum number.
8. *Run Simulation:*
  a. Run Simulation will generate a new set of *n* trajectories with the current settings including Simulation of speed, Arena, and Durations. You must click this button after changing any simulation parameters for the changes to take effect and appear on the trajectory map. This will erase the current trajectories and cannot be undone, so be sure that you have exported your previous trajectories before clicking on this.
  b. This is automatically run when the program first opens, you do not have to manually click it every time you open it
  c. This is used to generate multiple sets of trajectories. Rather than closing and reopening the program, you can just click on this button to start a new batch of trajectories
9. *Arena Sizing and Scaling*
  a. Bounded mode: Set the trajectory generation mode to Bounded to generate smooth, curved trajectories based on the specific angular and linear velocities while keeping the target within the defined arena boundaries.
  b. *Arena size:* Customize the size of the arena bounds an x and y height and width, represented by the entire border of the trajectory simulation window
  c. *Arena Scaling:* The proportion of the area that bounds the randomized path, defaulted to 0.88 for x and y. This is represented by the dashed line in the simulation window
  d. *Wall Avoidance:* Controls the tendency for the generated trajectory to stay a certain distance away from the walls when controlling its path. These overall control the spread and shape of trajectories
    i. *Wall buffer:* Determines the distance from the wall, in centimeters, that it will avoid running near.
    ii. *Wall Nudge*: Determines the strength of a pull to the center of the arena, and thus further away from the walls.
    iii. *Nudge Max:* How much the wall nudge pull is allowed to pull the heading direction of the dot per frame.
  e. *Starting Position:* This is used to control the tendency for where the dot starts is trajectory
    i. *Start Grid:* Determines the grid dimension, which allows for different numbers of possible starting positions.
    ii. *Edit Grid*: This will show the grid of possible starting positions, and which are activated (green) or deactivated (red). Clicking on a grid square will change it’s setting. The dot will then be generated to start in those areas.

*Other Parameters*.

1. *Angular Distribution*: changes the shape of the <u>angular probability distribution</u> (bottom right) that is sampled to change the angle of the trajectory. Options include:
  a. Gaussian: Favors tuning angles near the center of the selected angular range. This biases the trajectories to move in straightward directions.
  b. Wrapped: peaks at extreme angular velocities, set in the parameters column. This biases trajectories to have many clockwise and counterclockwise turns during the target trajectory.
  c. Skewed: quickly hit the peak towards the left side of the distribution and slowly go back down as it approaches the right side of the distribution
  d. *Angular Min*: extreme counterclockwise side the angular distribution (0°)
  e. *Angular Max*: extreme clockwise side of the angular distribution (60°)
  f. *Gamma Peak*: peak of Skewed distributions; actually “mu” in mathematical terms, but called “gamma” b/c Skewed is actually called a gamma distribution.
  g. *Kappa*: steepness of the peaks for <u>Gaussian and Wrapped</u> distributions.
    i. Higher kappa = thinner peaks (i.e. peaks are more “strict”, and will frequently only sample a few numbers).
  h. *Angular Shape*: steepness of peaks for <u>Skewed</u>.
    i. Alpha value for the Skewed/Gamma distribution.
  i. *Angular Scale*: speed at which distribution falls from the peak as x-axis value increases.
    i. Higher Angular Scale = slower falling.
    ii. The mathematical term for this is theta
2. *Linear Distribution*: changes the shape of the <u>linear probability distribution</u> that is sampled to change the speed of the trajectories
  i. *Linear Min*: minimum speed (cm/s) of the linear distribution
  ii. *Linear Max*: maximum speed (cm/s) of the linear distribution

*Trajectory Parameters*.

1. *Operations per Second* (HZ): frame rate of the camera in use, defines how many coordinates are sampled.
2. *Turn Min and Max: Sets the minimum and maximum number of frames between updates to the target’s speed and tuning angle. Smaller Turn values produce more frequent changes in movement resulting in tighter curves or smaller loops in Gaussian and Skewed modes. Larger values allow the target to travel farther between updates producing broader curves or larger loops*.
  a. *Note: Turn setup can strongly influence trajectory shape, even when the angular and speed distributions remain unchanged*.
3. *Operations per Trajectory*: how often the linear and angular vectors get sampled and concatenated (e.g. every x number of coordinates). If set to your camera frame rate a new vector is sampled every second. Setting this above or below the camera frame rate will generate more simple or complex trajectories, respectively.
4. *Occlusion:* Use this parameter to make the dot temporarily disappear either via ‘masking’ or insertion of an object.
  a. Gaps: Set the total space of the trajectory you want to have be invisible, and where it begins.
  b. Choose the position and size of the “object” that will block the dot during its trajectory.
5. *Duration*: temporal duration in seconds of each trial individually.
6. *Extend*: trajectory starts going in reverse for whatever the amount of time is designated
7. *Trim*: shortens trial by however long is indicated.

### A BONSAI-BASED WORKFLOW FOR TARGET PRESENTATION AND SYNCHRONIZATION

Bonsai is an open-source, visual programming environment that specializes in efficient processing of inputs and outputs to interface with external components connected to the software’s host system^17^. The versatility of the software and its expansive library of extensions permits control of video recording functionality, live behavioral tracking, trajectory rendering, and interactions with auxiliary software. In our apparatus, Bonsai serves as the central control system, coordinating five key functions: 1) visual target presentation and projection calibration, 2) real-time tracking of mouse and target positions, 3) target interception detection and trial control, 4) automated start chamber operation and reward delivery through Arduino and BPod interfaces, and 5) video acquisition and storage of behavioral data in CSV files. The script is accessible and easy for daily use by behaviorists. There is a template script attached in the public GitHub repository found for this paper.

#### Computer and Hardware requirements

The behavioral system was operated on a Windows 11 PC equipped with an Intel 14700KF CPU, NVIDIA 4070Ti Super GPU and VRAM, and 64 GB RAM capacity. These are above the minimum required specifications, but the system must support simultaneous operation of Bonsai, DeepLabCut, and MATLAB with Bpod, with sufficient GPU performance for real time pose estimation at 30 Hz. A solid state drive (SSD) is required for high speed video recording, along with a high bandwidth USB interface for camera acquisition. Detailed hardware specifications, software versions, and the system configurations used in our setup are provided in the **Materials List (Table 1)**.

#### Bonsai Set-Up

Download and install the latest compatible version of Bonsai for Windows from the official website (https://bonsai-rx.org/docs/articles/installation.html). To configure Bonsai for our behavioral apparatus, download the required files from the `Bonsai` folder in the accompanying GitHub repository (https://github.com/alab-ucsb/alab-pursuitRig). This folder contains two important files `bonsai.config` and PursuitBonsaiTEMPLATE.bonsai, along with four image files (`greendot.png, forest-green-sleep.png, f3sprite_t.png, obstacle.png`) used for visual stimulus presentation.

First, place `bonsai.config` in the Bonsai configuration directory, replacing a blank config file (file path is likely “`C:\Users\user\AppData\Local\Bonsai\Bonsai.config`”). This file specifies the packages required to run the workflow and its associated functions. You will likely need to update packages through `Tools > Manage` Packages after launching Bonsai.

Next, place `PursuitBonsaiTEMPLATE.bonsai` and four image files in the directory where daily recording folders will be saved. These will be read by Bonsai in order to load the images onto the canvas.

This is our customized Bonsai workflow for pursuit tasks. The main workflow canvas, along with the `Toolbox` and `Explorer` panels located in the sidebars. In the highest group, you will find the main workflow. This acts as the “front end” where session types are activated, trajectories are chosen, and session details are edited. A screenshot of the workflow interface is shown in **Figure 3a**. Using this template, new features can be added to adapt to your individual needs, and new session types can be added. Much can be discovered by exploring this template, and numerous features can be extended from the basic workflow.

**Figure 3.**
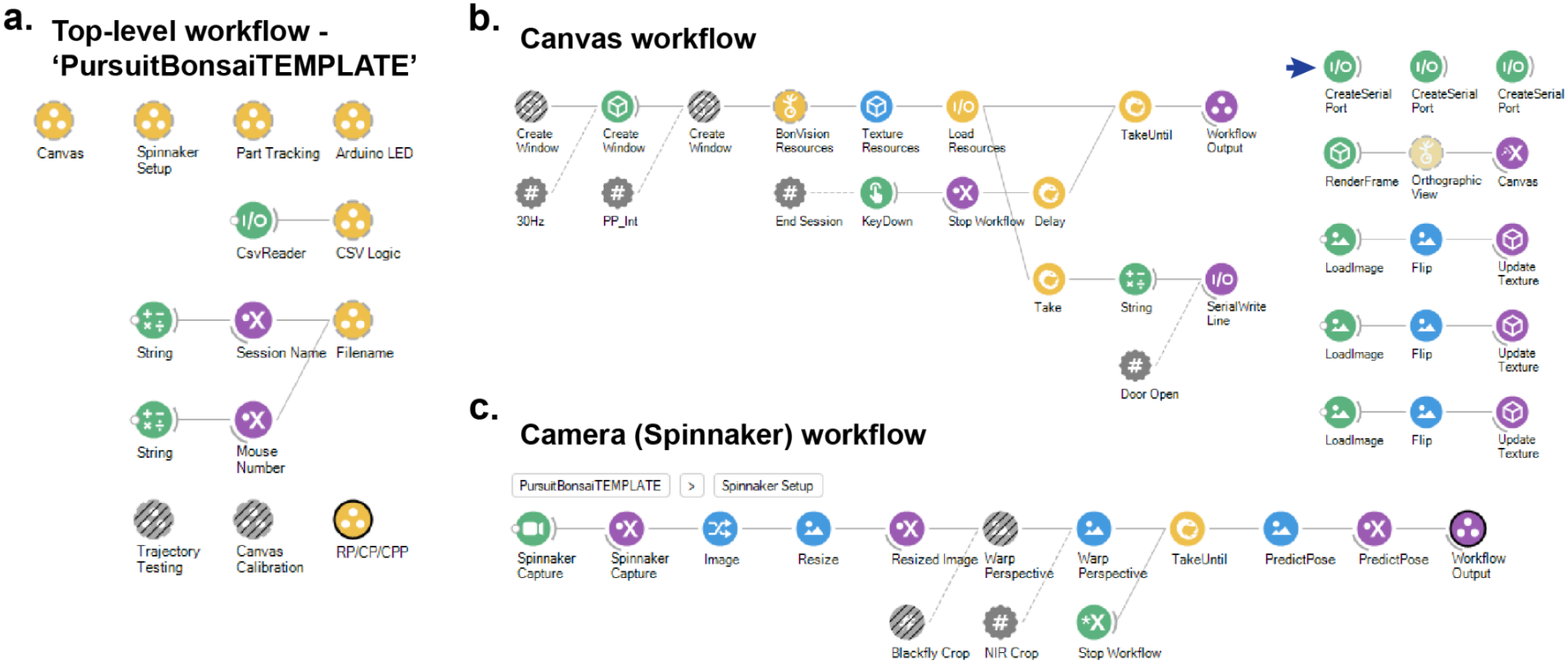
Bonsai workflows for the automated pursuit system. **a.** Top-level workflow (‘PursuitBonsaiTemplate’) can be downloaded from our Github repository and is used to control target chasing experiments after hardware setup. See **Protocol 3** for how to change node parameters for your system, **b**. Canvas workflow (‘Canvas’) for projecting the moving stimulus and changing COM ports related to automated reward delivery, start box, and TTL synchronization, c. Camera workflow (in our case Spinnaker) for viewing real-time position tracking inside the behavioral arena. Nodes with a blue arrow indicate locations where COM ports may need adjustment when building a new system.

#### Canvas Creation

In order to enable projection of pursuit stimuli, you will first need to create the canvas using the `Canvas` group in the Bonsai workflow. This group contains nodes for serial COM port configuration, rendering projection elements, loading images used in trials (including the green dot), loading external resources, and terminating the workflow using the “ctrl+space” keyboard shortcut. Serial communication is managed through serial read nodes (`SerialRead` and `SerialReadLine`), which receive data from external devices, and serial write nodes (`SerialWrite` and `Serial WriteLine`), which send data or commands to these devices (**Figures 3**,**4**).

In the provided template, COM3 is assigned to the Bpod State Module, COM7 to the Arduino responsible for irregular LED TTL signals and camera synchronization pulses, and COM8 to the Arduino controlling the automated start chamber. Because COM port numbers are assigned by Windows and may vary across systems, first identify the port associated with each device in Windows `Device Manager` under `Ports`. In Bonsai, select each corresponding serial communication node and choose the appropriate COM port from the `Port` dropdown menu.

Update the COM port assignments in the following locations **(Figure 4, blue arrows):** (1) In the `Arduino LED` group: Set the port corresponding to the TTL and camera synchronization Arduino (COM7 in our setup) in the `SerialWriteLine` node in the top workflow sequence (starting with `String`), the `SerialReadLine` node in the middle sequence, and the second `SerialWriteLine` node in the bottom sequence (starting with `Timer`). (2) `RP` group: set the Bpod State Module port (COM3) in the `SerialRead` node and the automated start chamber Arduino port (COM8) in `SerialReadLine` and `SerialWriteLine` nodes. Last, Within the `RP` group, locate the `SelectMany` node, open its workflow sequence and set the `SerialWite` node to the Bpod State port (COM3).

**Figure 4.**
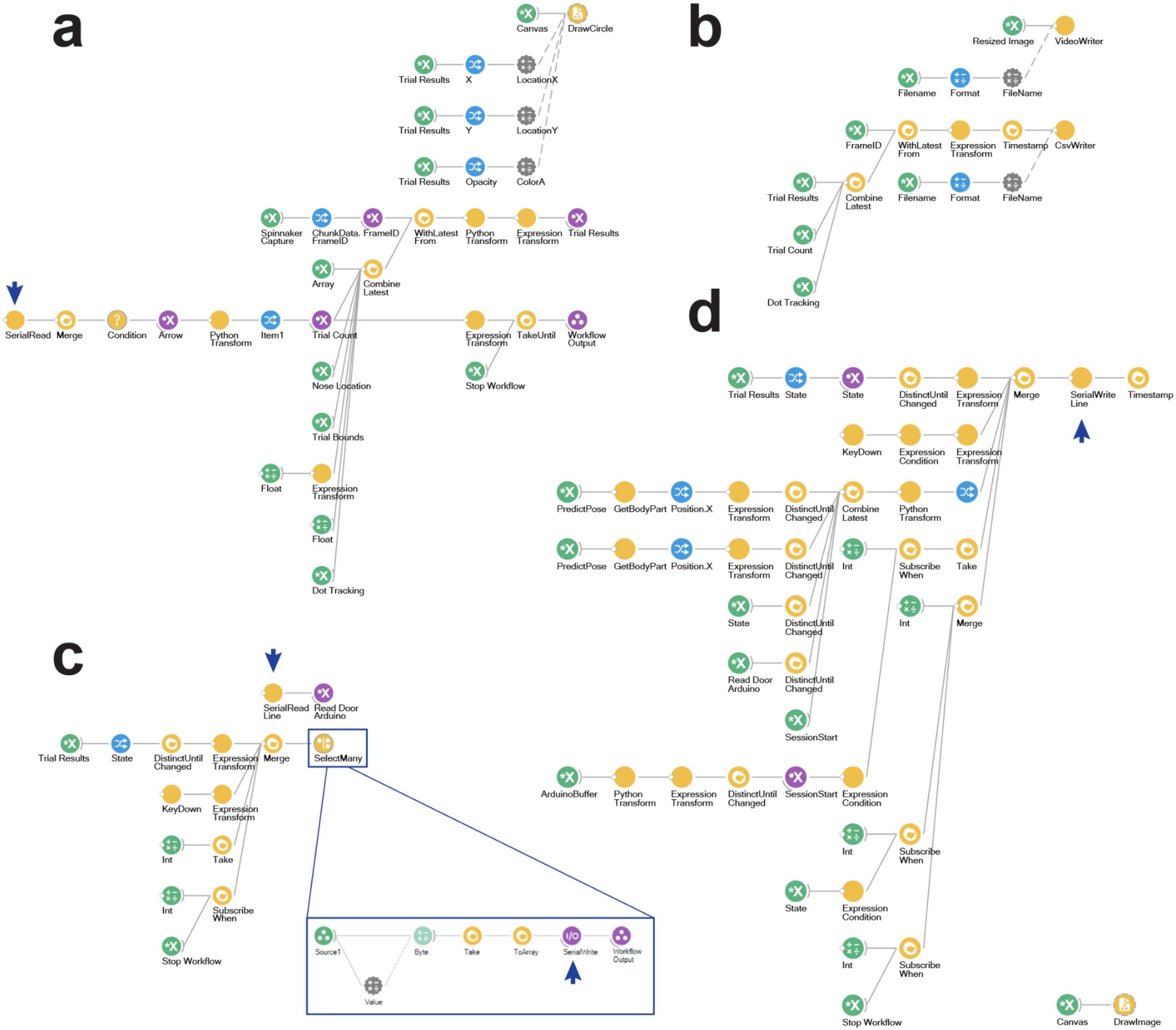
Nodes for Setting Serial Ports in the RP workflow. **a.** Trial and prediction logic, **b**. CSV and video writing, c. Trial control, **d**. Live tracking and door control. In the full workflow, **a-d** are arranged in sequential vertical order but are presented in clusters here for visualization. Nodes marked with a blue arrow are those that may require a change of COM ports. Each will default with COM3, C0M7, or C0M8 and must be swapped with your corresponding system’s serial port (Bpod, TTL Arduino, Door Arduino respectively).

After configuring the serial inputs, verify the visual stimulus settings in the remaining `Canvas` group. The Bonvision package handles the drawing of motion and other visual stimuli. In order to show the images, you will need to point the node to the correct `.png` in the `LoadImage` nodes. Examples are provided in the repository.

#### Camera Setup and live animal tracking with DeepLabCut

Animal behavior is recorded and processed in real time during target pursuit using a machine vision camera mounted above the arena. We use a Spinnaker Teledyne FLIR camera. Other brands of machine vision cameras (Basler, Allied Vision, etc.) are compatible with our system. Camera field of view needs to be able to fully capture the arena and startbox. This guide will focus on Spinnaker cameras. The camera is operated using Bonsai and the Bonsai workflow for setting and running up the camera is included in **Protocol 2. Figure 5** depicts electronic wiring for triggering and synchronization of the camera with the remainder of the system. For a Teledyne camera, the `Spinnaker Capture` node is used but a different capture node will be required depending on manufacturer (more details on Bonsai and nodes found in **Protocols 2**,**3**). To properly configure your camera in Bonsai check the documentation for your respective capture node.

**Figure 5.**
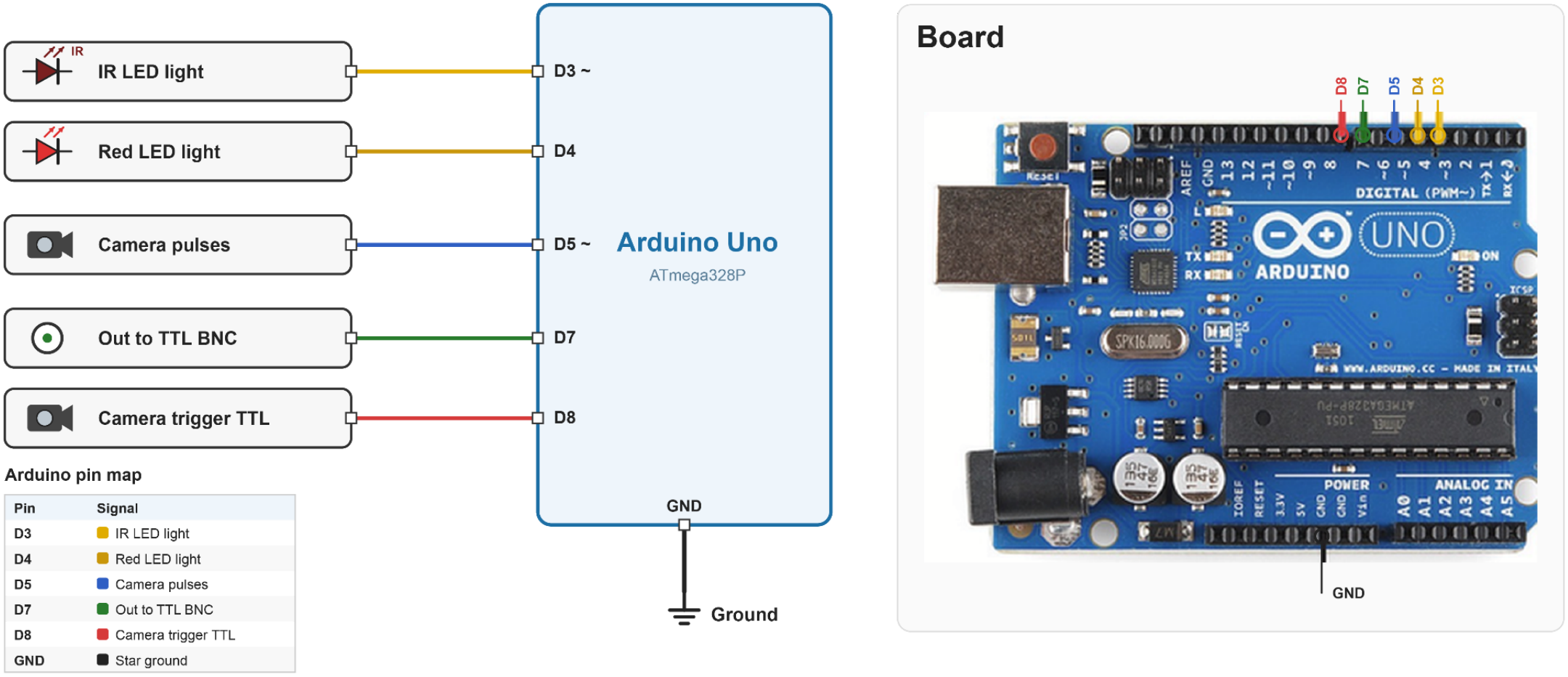
Camera triggering. Schematic diagram depicting connections between machine vision camera and Arduino for synchronization with an external clock or electrophysiology recording system (may differ depending on specific camera).

DeepLabCut (DLC) is an open-source deep-learning pose estimation software used primarily for animal tracking^18^. This software allows users to import videos and mark body parts in frames for DLC to learn. The final product is a model that can be used in DLC’s analysis functions and can additionally be exported for external software to identify body parts in videos. DLC offers a Bonsai library with modules for live behavioral tracking with low latency, which requires one of these models^19^. The current `Bonsai.DeepLabCut` packages requires TensorFlow based models and does not natively support PyTorch trained models.

We trained a DLC model using videos from trial experiments with our behavioral setup to monitor for nose, right ear, left ear, tail, and target location. Accurate tracking was essential for reliable target interception detection and automated door control. To improve model robustness, we recommend including training frames that capture variations in environmental conditions, animal positions, and body orientations. Model performance should be validated across representative behavioral conditions. Our trained model achieved a training error of 1.83 pixels and a test error of 3.23 pixels, consistent with the tracking accuracy reported in previous DLC studies^18,20^. SLEAP can also be used for live tracking and has a Bonsai package that can be easily swapped into the Bonsai workflows provided with this protocol^21^ (https://github.com/talmolab/sleap).

In some cases, optimizing settings related to the capture of the camera requires modifications in the corresponding Software Development Kit (SDK) provided by the manufacturer. In the Bonsai workflow, we use a `SpinnakerCapture` module that we modified to be compatible with our camera to stream a live feed at 30 Hz from the camera at a defined ROI. To optimize the processing speed, we also resize the frames from 1224 px x 1024 px to 640 px x 480 px before outputting them to a `PredictPose` module from the DLC library. This module processes each frame of the video and outputs a coordinate for any visible parts, using the resized camera frame as a reference coordinate plane (i.e. the x-axis spans [0, 640] and the y-axis spans [0, 480]). The coordinates are then rescaled and normalized to a range of -1 to 1 to be output to a Bonsai canvas that gets projected to the Open Field Maze floor. For calibration purposes, an optional visual marker displayed the tracked nose position on projected canvas, allowing users to verify tracking alignment. This marker could be enabled or disabled by setting a parameter to 1 or 0, respectively, and remained hidden during the behavioral experiment. This setup yields continuous position tracking for online target interception detection. The tracked coordinates and corresponding timestamps were simultaneously saved to a CSV file during each behavioral experiment for data analysis purposes.

#### Position Prediction and Capture Logic

There is latency between the projection system, camera, and real-time tracking that makes simple capture logic between the dot and nose inadequate. To compensate for this, we have created a system of predicting the real-time position, velocity and heading of the animal from prior position estimates which is used to trigger captures. Using Bonsai, a script (“`Pred Trial Logic`”, inside `RP` node) calculates the animal’s velocity using the difference between the latest DLC position output of the nose and the position of the nose 5 frames before, then dividing by 5 (number of cached frames) for both x and y. A lookahead prediction of the nose’s current position is calculated using the latest DLC position plus two times that velocity. This prediction is the predicted current position of the nose, which we use in order to compensate for the latency between the systems. This prediction is then used to calculate the distance between the dot and the animal’s position. The head direction vector of the animal and the angle to the dot is calculated simultaneously. Position of the moving dot is derived from the input CSV drawing the dot, rather than the tracked DLC position. This is done for the same latency compensation reason. The predicted position and head direction are then used to gate a successful capture by combining a capture radius of 0.1 cm and a capture angle sector of ±35° from the animal’s heading direction (these values can be customized).

#### Bpod, States, and Trial Logic

The target chasing behavioral apparatus is fully automated through Bonsai which, as described above, tracks stimulus capture in real-time and controls several hardware components that impose trial structure and deliver water reward (**Protocol 3**). Water is dispensed following target interception using a Bpod state machine, valve module system, and a behavioral port featuring integrated nose port detection and water delivery (Bpod, Sanworks). The Bpod valve is mounted inside the start chamber at an appropriate height for the animal (discussed in the automated start box section). Nose pokes are sensed by the valve and water is only released after successful trials in which the animal intercepted the visual target. The amount of water to be released by the valve is controlled by the Bpod GUI run by MATLAB. The Bpod water delivery system should be calibrated before initial use and recalibrated whenever the height of the water delivery apparatus is adjusted to ensure accurate reward volumes.To activate Bpod state control run `Pursuit_Protocols.m` in MATLAB (available at https://github.com/alab-ucsb/alab-pursuitRig/tree/main). This script controls the activation of Bpod, a graphical user interface (GUI), and writes an output spreadsheet containing a state, output or event, with an accompanying state name and timestamp. Parameters define the amount of water that is released by the port and when its signaling. This script will need to be run, just once at the beginning of experimentation, to activate Bpod and connect to Bonsai which runs the remainder of the processes for closed-loop target chasing. While the Bpod script is straightforward, for any troubleshooting refer to the Bpod documentation^22^ (https://sanworks.github.io/Bpod_Wiki/).

When running the Bonsai workflow for this system (**Protocol 3**), there is an organization of states that control the progression of trials in a session. The Bpod module is capable of sending and receiving bytes in order to signal changes. We have implemented 6 different states to work in tandem with these byte signals (sent by Bonsai to Bpod) to change trials, and document successful captures and session events. State 1 is the system waiting at the beginning of the trial with a visible and stationary dot (e.g. animal in the start chamber after nose poking). State 2 is when the dot is moving (i.e. a chase state). State 3 is a successful capture (lasting 1 frame before changing). State 4 is a timed out trial/missed capture (lasting 1 frame before changing). State 5 is a holdover until the next nose poke of the Bpod, which initiates the next trial. State 6 indicates that the session has completed. These states control trial automation and run consecutively with only input from the animal. Bonsai sends bytes to Bpod to indicate trail checkpoints, and Bpod sends a byte 12 when the nose poke is registered which increments the trial.

### Running Pursuit

As a layout for the most common session type, there is an `RP` node that is used to run experimentation (this must be the only node activated in the bottom row of the main workflow). This is used to create a trial by trial automated session. This automates the pursuit stimuli, trial iteration, and water delivery (discussed further below). Additionally, it handles the capture logic during experimentation.

The `PursuitBonsaiTEMPLATE` top-level workflow controls the full experiment. The workflows are presented in **Figure 4** and explained step-by-step in **Protocol 3**. After activating the desired node on the bottom, change the `String` values for animal number and session type, then load in the file path to the trajectory CSV you’ve created (See **Protocol 1**). Hit `Run` in order to run the workflow, and it will begin, canvas should begin calibration, and LED startup flashes start. When running for the first time, it is useful to see the recording viewer and tracking render. To enable a visual of what the camera is recording, navigate to the `Spinnaker Setup` group. Then, double-click on the `Image` and `PredictPose` nodes.

The `Filename` group in the top-level workflow holds logic for autosaving results of a session. This workflow automatically takes parameters from the user to name the file according to the animal and session type. This will then automatically save a folder for the animal, date and session where data CSVs and video are saved. A Python node in this workflow also saves the results of the Matlab file running Bpod (discussed further below). In order to change the saving location, you will need to enter this group workflow (`Filename`) and replace the starting string in the expression script node titled `Format Filename`. It will default as a drive string that can be replaced with any folder location path, and it will fill-in a date folder, animal folder and populate it with generated CSVs and video.

After any recording session is finished, it is crucial that **ending the workflow is not done by pressing the stop button. Instead, end the experiment by pressing “Ctrl+Space”**. This will end recording for all systems at the same time, as opposed to just the Bonsai workflow.

**Protocol 2 - Wiring of Camera Pulse and Irregular TTL for synchronization**.

The wiring diagram for this microcontroller is displayed in **Figure 5**. This will control the camera trigger pulse and an irregular LED TTL pulse that occurs every 3-15 camera pulses that is used as a backup for synchronization in case there are dropped frames from the camera.

1. Connect the Arduino microcontroller to the computer using a USB Type-B cable.
2. Use a perfboard to connect the Arduino to an external ground (see in *Electrical Architecture and Grounding*).
3. The ground wire of all BNC cables and LED circuits are connected together using a star ground connection point.
4. Connect the signal wires of BNC cables and LEDs to the corresponding Arduino pins shown in Figure 3. These pin assignments correspond to the `randomLEDTTL` code provided in this rig’s code repository (https://github.com/alab-ucsb/alab-pursuitRig).
5. Power the LED lights using the microcontroller’s 3.3V supply, with a 100Ω resistor connected in series.
6. IR and colored LED lights can be run and affixed taped to the side of the arena, ensuring that their flashes are detectable by the camera.
7. Machine vision cameras typically have a general purpose input/output (GPIO) interface that can be used to trigger image acquisition or transmit data from the camera to external clocks. For the Teledyne setup described here, there is a circular Hirose 6-pin connector and corresponding cable. This cable is run from the camera port and has 6 colored wires that can pass current and be configured for different inputs or outputs.
  a. Refer to the pinout documentation for the specific camera cables. In our setup, the red wire is used for the signal connection and the brown wire for ground.
  b. The red wire can be run back to the microcontroller, and connected to pin 5.
  c. The ground should be run to the star ground we started earlier.
  d. The remaining wires can be capped to reduce electrical noise and prevent short circuits.
8. After establishing the ground connections, route the BNC cables from the microcontroller to the BNC breakout board connected to your acquisition system.

*Camera Trigger Activation for Teledyne Cameras*.

To pass TTL pulses from the camera, you will need to configure particular input/output lines in the SpinView application.

1. Open SpinView and connect to your camera
2. Navigate to the Features panel on the right side and expand the Digital IO Control (or the GPIO tab).
3. Expand Acquisition Control and change the following parameters
  a. Trigger Mode: Set to On.
  b. Trigger Source: Select the exact line you wired your pin to (e.g., Line 0 if you used Pin 2, or Line 3/Line 2 depending on your model).
  c. Trigger Selector: FrameStart.
  d. Trigger Activation: Set to Rising Edge or Falling Edge depending on how your external device builds its signal

**Protocol 3 - Running Pursuit in Bonsai**

1. Launch the Bonsai application, and open the downloaded `PursuitBonsaiTEMPLATE.bonsai` workflow.
2. The main workflow serves as the user interface (**Figure 4a**). The top row contains four groups (`Canvas, Spinnaker Setup, Part Tracking,` and `Arduino LED`) for configuring projection parameters, camera acquisition, DLC tracking and Arduino communication. The bottom row contains three experimental modes: `Trajectory Testing` for visualizing GUI generated trajectories, `Canvas Calibration` for adjusting the projection canvas, and `RP` for running pursuit experiments. Additional nodes can be created as needed.
3. To run the randomized pursuit task, enable `RP` node and disable `Trajectory Testing` and `Canvas Calibration`.
4. Specify the trajectory file by double-clicking on the green `CsvReader` node (**Figure 4a**). Replace the existing file path with the path to randomized trajectory CSV file generated in **Protocol 1**.
5. Configure the output file names by editing `String` nodes corresponding to the animal identifier and session name (**Figure 4a**). The session data is saved automatically.
6. Configure the COM ports before running the workflow for the first time.
  a. Double-click on the yellow `Canvas` node (**Figure 4a**), this will open the ‘Canvas’ workflow (**Figure 4b**)
  b. Locate the three green `CreateSerialPort` nodes at the top of the workflow (upper right **Figure 4b**)
  c. Change them according to the identified serial ports on your computer, where these are the default ports in the template: COM3 is the Bpod State Module, COM7 is the irregular TTL + camera pulse Arduino, and COM8 is the Arduino controlling the automatic start chamber (**Figure 4b**)
  d. The COM port will need to be changed in the following nodes as well (replace with your corresponding port number): (1) `Arduino LED` group (**Figure 4a, upper right**) → COM7 in `SerialWriteLine`, `SerialReadLine`, `SerialWriteLine`. (2) `RP` group (**Figure 4b, lower right**) → COM3 in `SerialRead` and COM8 in `SerialReadLine`, `SerialWriteLine`. Refer **Figure 4c** to locate the `SelectMany` node within the `RP` group → set the `SerialWite` node to the Bpod State port (COM3).
7. Click **Start** in Bonsai to execute the workflow. Two visualization windows should appear: one displaying live video of the chamber and animal, and the other showing the animal in the open arena with annotated DLC tracking results. If the windows do not appear automatically, open `Spinnaker Setup` sequence and double-click `Image` and `PredictPose` nodes to display the camera acquisition and DLC tracking outputs, respectively, while the workflow is running (**Figure 4c**)
8. To terminate the session, it is crucial to press **Ctrl + Space** on the keyboard to stop all experimental systems simultaneously. Do not use the STOP button in the Bonsai interface, as this does not properly terminate the irregular LED synchronization and the TTL signal generation.

#### Synchronization methods

In order to synchronize the different systems and softwares that run in this rig, we have introduced a system of Transistor-Transistor Logic (TTL) pulses from hardware modules onto a centralized clock. While this could be accomplished using a standard data acquisition device (e.g. a National Instruments DAQ), in our setup we pass TTLs into an IMEC OneBox (IMEC) run with OpenEphys Acquisition Software^23^. In either setup, TTLs passed onto a common clock can be used to establish common reference points for Bonsai, Bpod, and Arduino to synchronize behavior (and neural) data.

TTL pulses are programmed to trigger in a structured manner depending on their source. All TTL pulses are wired to a BNC breakout board connected to our OneBox but any other input/output interface would work similarly. Using a Bpod State Module, a 1-second pulse is sent when water is delivered from the port and a 0.5-second pulse is sent when trial success signals are received. Using a dedicated Arduino Uno microcontroller board, 30Hz camera trigger pulses are received and wired to the camera’s state system to fire in sync with the video capture frame-by-frame. Tied to these pulses are an irregular pulse that triggers at random intervals between 3 and 15 camera pulses. These pulses control an IR LED that is visible within the field-of-view of the camera. They always fall on a camera pulse, and provide an irregular reference pulse chain. Using another Arduino microcontroller that controls the automated start chamber (described below), a single pulse is sent whenever the door is opened, and two consecutive pulses are sent when the door is closed. In order to define a starting point, a recording starts with 5 seconds of 5 Hz pulses then 3 pulses triggered by the door microcontroller for the start of the session. This acts as a session starting point for data analysis and training stimulus for the animals. All pulses are wired to and recorded by the common clock, and individual CSVs are written by Bonsai depending on which module they come from. Bridging these pulses with custom software allows for data synchronization, recovery and analysis.

### CONSTRUCTING THE PHYSICAL TARGET CHASING RIG

#### Automated Start Chamber

A key feature of our system is an automated start chamber that integrates trial progression and reward delivery to support repeated behavioral testing with minimal experimenter intervention (**Figure 6**). The start chamber (30.5 cm x 23 cm x 28 cm) is affixed outside of the pursuit arena. A horizontally sliding door separates the chamber from the chasing arena, controlling the animal’s entry into and return from the arena during each trial. This design minimized the distance between the chamber and arena while accommodating mice with or without neural recording tethers, as the horizontal door movement avoids interference with the recording cables. The chamber serves as the starting and ending point of each trial, as well as the location where a water reward is dispensed after successful trials (see next section). All CAD files for building the chamber are available in our GitHub repository (https://github.com/alab-ucsb/alab-pursuitRig)

**Figure 6.**
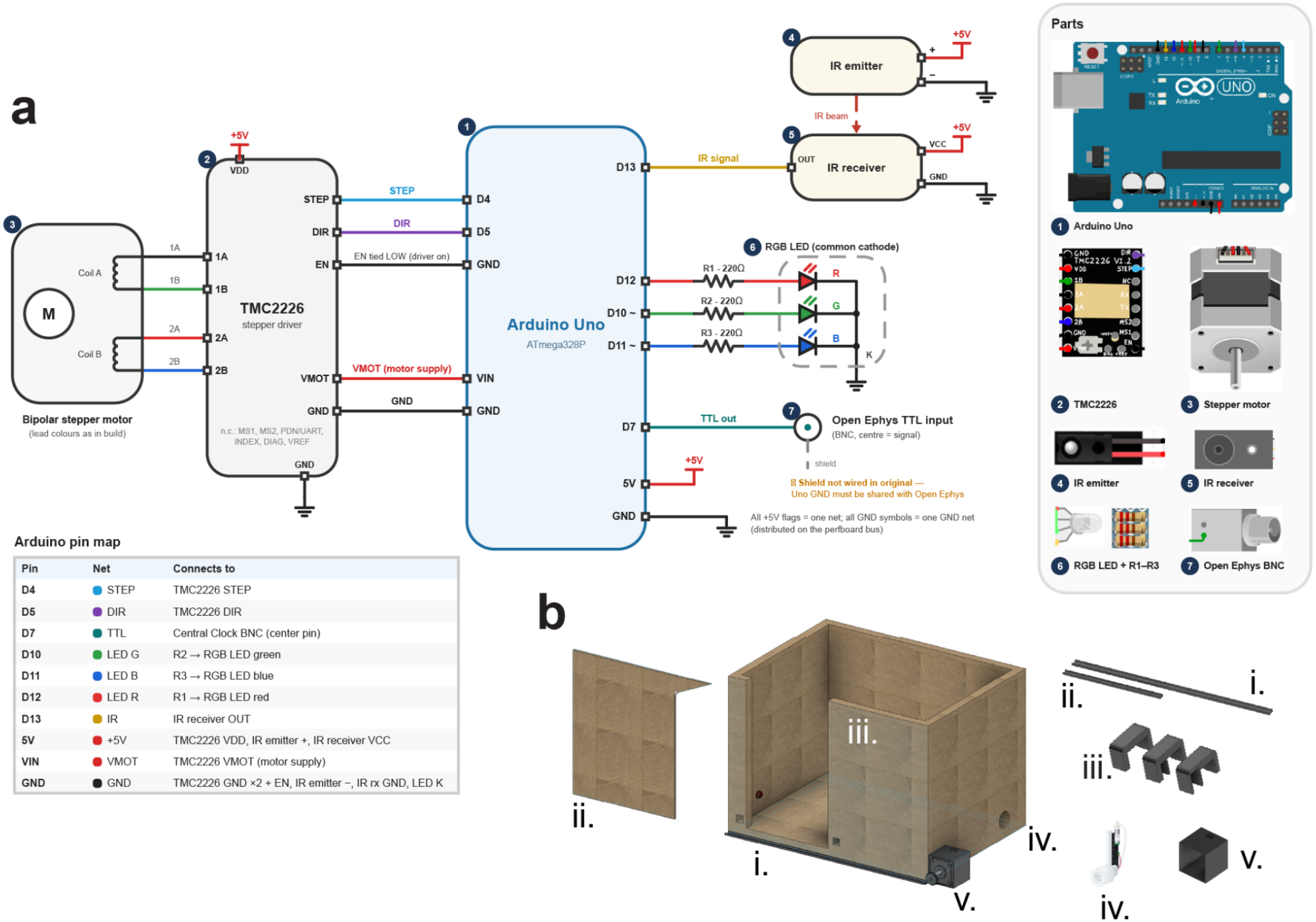
Automated starting chamber. **a**. Wiring schematic for stepper motor, driver, infrared door trigger, Bpod water delivery, and LED for synchronization. Left diagram, schematic wiring diagram. Right column, required components. Bottom left, table of connections, **b**. Diagram for assembling the starting chamber. Assembled box on left, 3D printed components on right. Numbered labeling indicates approximate location on starting chamber that 3D printed components are attached to.

**Figure 7.**
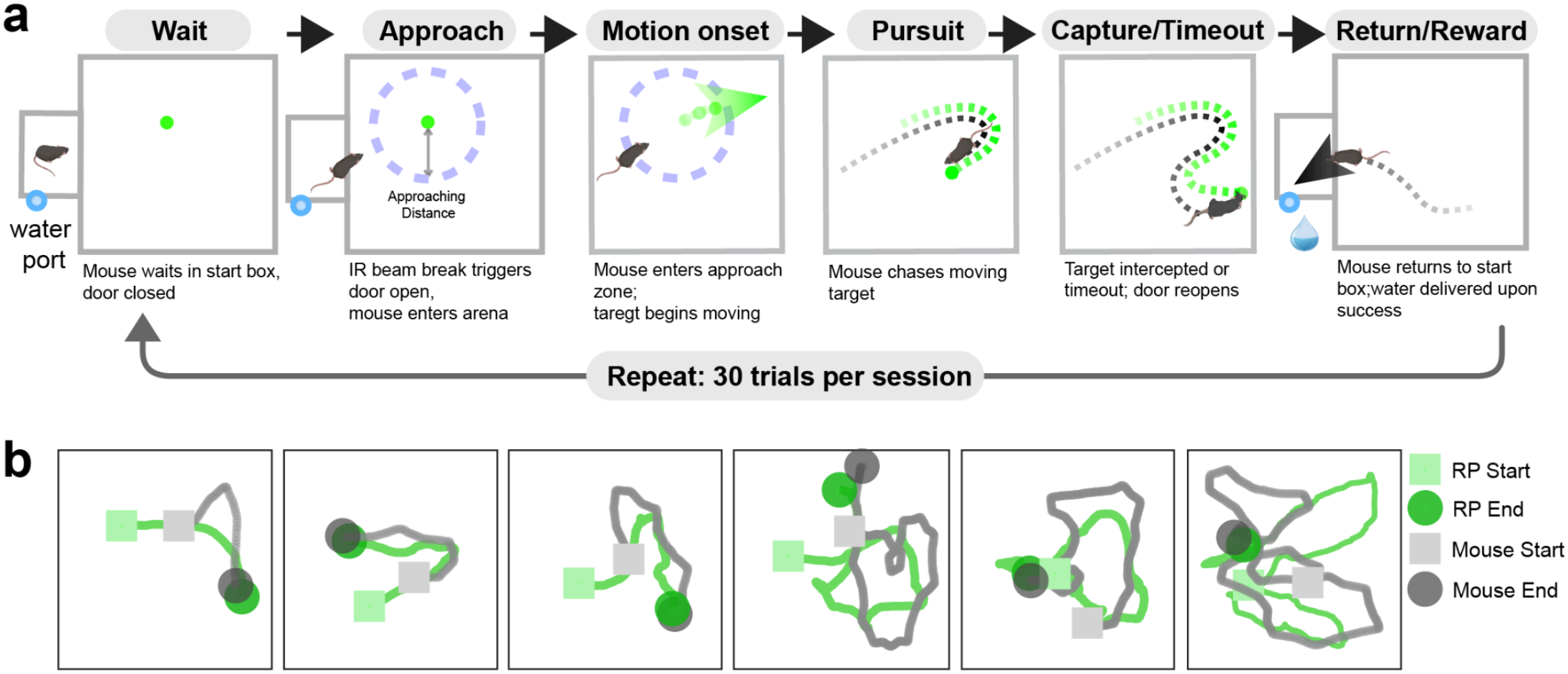
Schematic of pursuit trial phases and example chasing bouts. **a**. Structure for an individual automated pursuit trial. Mice wait in the automated start box and trigger the trial by breaking an IR beam near the door. The dot is positioned in a random x,y-coordinate which the mouse must search for. Once the mouse enters an approach distance within ∼30 cms diameter, the target begins moving according to the pregenerated movement trajectory. The animal will typically engage in pursuit until interception. If the animal is unsuccesful, the trial times out which typically is set to ∼10s in our experiments (i.e. a failure trial). After trial timeout or successful interception the animal returns to the automated start box to wait or retrieves a water reward from the Bpod port, respectively, **b**. Six example pursuit trials with the mouse in gray and the target in green. The start is indicated by squares and the trial end is demarcated by circles.

This start chamber is integrated with the Bonsai workflow to automate trial progression based on the animal’s behavior. The door can be controlled automatically using infrared (IR) beam detection and animal position-based tracking with DeepLabCut, or manually using physical buttons or keyboard commands through Bonsai. Door movement is controlled by a bipolar, NEMA 17 stepper motor driven by a TMC2209 stepper motor driver (Adafruit) operating in StealthChop mode for precise, consistent, and quiet operation. The stepper motor provides sufficient torque to reliably move the door without stalling. **Protocol 4 and Figure 6** describe how to construct and connect the start chamber for control via Bonsai and animal behavior.

**Protocol 4 - Constructing the automated start chamber. (Figure 6)**

*Construction of the start chamber*.

1. Saw or lasercut ½-inch (1.27 cm) plywood or other material into proper sizes. Give attention to making straight cuts, and level edges to promote an angular box (Figure 5).
2. With properly sized pieces, saw the doorway into the front wall, drill a 2.3 cm circular hole for Bpod on the right wall (with doorway facing towards you), and drill small holes for IR sensors to be run through. (Figure 5d). *Note:* The doorway can be cut before or after construction of box, but it is more difficult to saw the piece after it is erected.
3. 3D print the motor box, rails, and the door stabilizer (Figure 5, i-v).
4. Glue the bottom outer rail to the front bottom of the box. Glue the top door stabilizer to the left side of the doorway, aligned with the top of the door panel. Glue the inner rail to the bottom of the door panel. This allows sliding of door on the outer rail attached to the box
5. Assemble the door: slide the slider onto the track and confirm smooth movement with no binding. Lubricate the rails if necessary.
6. Assemble stepper motor box.
  a. Create an opening on the top of the 3D-printed motor box to run wires through.
  b. Line inside of box with foam for sound dampening and stabilization
  c. Run motor wires through the hole on the top of the box, and pull through to insert stepper motor into the box with the axle facing out (should be secure, but can still move in and out a little, as well as rotate, Figure 5b,v)
  d. Cut 2 pieces of anti-vibration pads to size and glue to the bottom and left side (so it goes between motor box and wooden wall) of the motor box.
7. Mount the stepper motor box to the right wall at the correct height so the gear teeth mesh cleanly with the rack on the door.
8. Assemble door panel
  a. Cut another piece of anti-vibration pad, this time to cover the width of the door and mount the gear rack onto
  b. Assure that anti-vibration pad is flush with both edges of door panel
  c. Glue left side of gear rack to anti-vibration pad, and make sure the corners of both are flush together so they can make a right angle on door.
  d. Place the door panel in the rail section and test slide first. Then position door panel in the closed position and align the gear rack at the correct height so that it sits parallel with floor and grooves sit snuggly with motor pinion. Glue gear rack at this height and angle so it slides smoothly without lifting.
9. Glue top rail onto front wall and wrap around top of door panel to stabilize
10. Confirm again door slides smoothly by hand-turning motor and having it pull the gear rack and door (Can add rail lubricant dust if necessary) *Automated start chamber electronics*.
11. For the full RGB LED light, connect Red to pin 11, Green to pin 10, Blue to pin 12 (each through a 220Ω resistor), cathode to GND. Upload LED test code and confirm all three channels light independently. (Figure 5)
12. Create 5V and GND power rails on the breadboard. Connect the sensor signal wire to pin 13. Confirm LOW = beam blocked, HIGH = beam clear. IR indicator light on Arduino board will show beam status. (Figure 5)
13. TMC2209 stepper motor: The motor will have two coils, each with two wires (A&B), that will be connected to the stepper motor driver. Connect motor leads and driver to pins 4 (STEP), 5 (DIR), 6 (EN). Upload stepper test code. Disconnect motor power before changing any wiring. (Figure 5)
14. Run the combined main firmware and confirm all subsystems interact correctly before soldering.
15. Make sure all wires are run long enough, preferably to just a little longer than the perfect length to wrap around the box. Full wiring
16. Run the RGB LED and IR sensors into their wall openings. Confirm the door slides without catching any wires or glue.
17. Solder longer lead wires to the RGB LED (color-coded) and IR sensors. Twist wire pairs for cable management. Apply heat shrink tubing.
18. Create a grounding bridge for LED, IR sensors, and buttons
19. Reproduce the validated breadboard layout on the perfboard. Keep the Arduino, TMC2209, and passive components logically grouped.
20. After soldering, re-run the full firmware test including Bonsai serial commands before closing up the box.

#### Maintenance of the Automated Start Chamber

As is common with systems relying on 3D printed parts and point-to-point wiring, degradation of components is possible, but easily replaceable. For the start chamber to be reliable for daily use over long term experimentation, it is wise to conduct semi-regular maintenance on its parts. This mostly constitutes cleaning of the rail pieces, stabilization of wiring, and diagnostics on electronics.

The most common necessity of maintenance is the rail and guide that the door slides on. Work with animals and daily wear-and-tear causes weakening of 3D printed focal points and grime to hinder the doors ability to slide. This can be diagnosed by increased friction while sliding or jumping off of the motor. It can be avoided by cleaning the rail with a vacuum or cotton swab, or reapplying rail lubricant dust. In cases of extreme degradation, replacing the rail or door guide is the best course of action. Mice may also push against the door, causing it to become misaligned with the motor gear. In such cases, manually readjusting the door alignment can restore normal operation.

Using a generic Arduino microcontroller leaves the possibility of wiring becoming disconnected from the pins. Stabilizing wiring in a place that avoids exposure to movement and wrapping wires prevents disconnections. Electronics in this system (motor, driver, LEDs) can also see reduced effectiveness over time. Pay mind to their possible malfunction over time, and how their replacement can be carried out.

#### Physical Structure

The structural framework of the behavioral mouse rig is constructed using 4040 aluminum extrusions to provide a sturdy, modular assembly fit for alterations. The dimensions of our rig structure is 89 cm x 86.1 cm x 208.9 cm and depicted in **Figure 1a**, but your assembly may vary depending on needs and physical constraints. Within the physical frame, a shelf is constructed 76.2 cms from the floor and a 0.95cm Plexiglass layer (McMaster-Carr Part #8774K46) is cut and positioned to span the inner boundary. A white projection screen is stretched atop the Plexiglass, and a top layer of 0.3cm Plexiglass is placed on top of the screen (McMaster-Carr Part #8774K16). The screen should be stretched evenly between two Plexiglass layers to prevent wrinkles that could distort the projected stimulus. Clips can be used to secure the curtain and maintain a smooth projection surface.

The target stimulus is projected onto the arena floor using a DLP projector (Ultra Short Throw Projector, Elite Projector) affixed to the base of the rig. The projector has a 0:3:1 throw ratio, allowing it to be positioned very close to the rig while still casting a large image onto the screen. In our setup, the projector is mounted 87.5 cm from the center of the screen. It is affixed as an outrigger to a cross beam, which is 71.1 cm below the bottom of the screen, run between two of the legs of the rig, near the bottom. The outrigger is 44.5 cm center-to-center from the cross beam (**Figure 1a**). At this distance, the projector is tilted at a 15° angle, and projects solely onto the table screen. The exact positioning of the projector will depend on the dimensions of your final setup.

The pursuit arena (61 cm x 61 cm x 31.75 cm) is constructed from plywood as a standard openfield maze, with walls painted gray to provide a uniform visual environment. An 18cm wide doorway is cut into one wall of the arena to allow access to a start chamber where trials are initiated and reward is delivered. The arena is centered above the projection screen and directly below the behavioral tracking camera. The entire apparatus is surrounded by alternating black and white curtains that provide global visual cues while minimizing external visual distractions during behavioral testing.

Although the behavioral tracking camera is sensitive to IR light, additional illumination is required for reliable animal tracking. To improve image quality, IR LED strips are installed along the top of the pursuit arena to provide more uniform illumination. The lighting arrangement should minimize glare and electrical interference that could introduce noise into electrophysiological recordings.

#### Electrical Architecture and Grounding

An important final step (especially if pairing with in vivo electrophysiology) is ensuring that all components of the automated pursuit arena are properly grounded. Stray magnetic fields, induction and electrical signals can result from things like the IR lighting, camera and automated motor. In order to mitigate these effects we have implemented preventative maintenance to reduce all electrical noise. All Arduino, BPod and TTL wires are grounded to a central metal cylinder clipped to the outside beam of the physical rig. Each of the grounding wires were fitted with metal clips that were clipped to the cylinder. The cylinder is then run by a cable into the surge protector outlet using a ground plug adapter. The external area of the open field arena, near where the motor and start chamber electronics lie, is lined with copper tape to act as an EMF field. A wire is connected to the shielding from the ground train on the start chamber perfboard. This eliminates induction caused by the motor driver activating the motor. Collectively, this grounds all electronics to the same source.

For researchers integrating the behavioral apparatus with in vivo electrophysiological recordings, additional measures may be necessary to minimize electrical interference. In particular, the Bpod control system and IR LED illumination can introduce recording noise that is not fully resolved by the grounding and shielding procedures described above.

#### TRAINING RODENTS TO INTERCEPT MOVING TARGETS

Rodents exhibit natural interest in novel objects or slowly moving visual stimuli, but reliable pursuit and interception of floor projected stimuli for reward requires behavioral shaping. Acquisition of visual target pursuit behavior requires approximately 2-3 weeks of training with individual variability between animals.

During the first 3 days, water deprived mice were habituated to the pursuit arena for half an hour per day. Water was manually delivered through a BPod water port to familiarize the animals with the reward location. Following habituation, mice began initial training with a slowly moving visual target at 5cm/s for approximately two days. Each trial initially lasted 30 to 60 s to allow sufficient time for animals to explore and approach the target. The target began moving immediately when the mouse entered a predefined approach zone surrounding the target (30 cm). Successful interception was defined as the mouse approaching within 1 cm of the target center, after which the target disappeared and water reward was delivered upon return to the start chamber. Trials without successful interception terminated without reward. Most mice learned to pursue and intercept the slowly moving target within a single session with 30 trials. For animals that had difficulty acquiring this behavior, an optional one-day stationary target training session (1-2 sessions, 30 trials per session) could be introduced before moving target to facilitate association between target interception and water reward. In our experience, starting with a stationary target or directly introducing a slowly moving target resulted in similar acquisition of pursuit behavior.

Over the following one to two weeks, target speed was gradually increased from 5cm/s to progressively broader ranges of 5-10, 5-15, 5-20 eventually reaching a maximal speed of 35 cm/s. Target speed was generally increased daily based on individual performance, with additional training at each stage could be extended as needed. Once mice became more proficient at intercepting the moving target, trial duration was reduced to approximately 15 s to improve training efficiency.

During initial training, mice completed one session per day, consisting of 30 trials and lasting approximately 30 to 60 minutes. As target speed increased and pursuit performance improved, training frequency was gradually increased to two or more sessions per day, depending on experimental requirements. In our experiments, mice completed three sessions per day, with 30 trials per session and a trial duration of 11 s. At this stage, each session took approximately 10 to 15 minutes to complete. Water reward volume was adjusted to maintain task engagement while providing sufficient reinforcement. Mice weighing 20 to 25 g generally exhibited reduced task engagement after consuming approximately 500 µL of water, although this threshold varied across animals. Training continued until mice achieved a stable interception success rate of at least 80% (**Videos 2-5**). Throughout training, animals were maintained under controlled water access, with body weight monitored relative to baseline (approximately 80% of baseline body weight).

## DISCUSSION

The protocol presented here outlines an entire system for the automated study of rodent target pursuit behavior using precisely controlled visual stimuli, including the generation of trajectories, the construction of an automated start chamber for introduction of trial structure and presentation of reward following interception, a protocol for the construction of the physical floor projection rig, electronics for running all components and synchronization for later data analysis, and a shaping protocol for training rodents to intercept purely visual moving targets. All code, Bonsai workflows, wiring diagrams, and CAD files are either presented here or available in our publicly accessible Github repository.

While the current system is highly efficacious for studying pursuit behaviors over numerous, highly-controlled trials, there are some caveats that should be considered when implementing the current protocol. Specifically, there are inevitable issues with latency, variability, and system failure. We have taken precautions to minimize the effects of such errors but experimenters should not build this system with the goal of running animals fully unsupervised.

Of note, there are occurrences of “ghost captures” where the animal intercepts the target, but the capture is not registered by Bonsai. This has only presented itself in cases where rodents were moving at a high speed at the time of interception such that the target is “eclipsed” for a very short interval. This likely occurs because of latency between the real world capture and the camera processing speed (**Table 2**) or in cases in which frames are dropped. This similarly affects the capture angle, as the lookahead is not exactly the animal’s heading, so it registers the predicted head direction, as opposed to the ground truth. We suspect that this can be optimized by changing video capture parameters that process the packet transfer between the camera and computer. This is dependent on the type of camera used and corresponding data transfer parameters which should be customized on a per use basis. We recommend checking the documentation of your camera for optimizing its transfer speed. One way to both reduce latency and dropped frames would be to modify our protocol to include a motion capture system. Common motion capture systems typically have below 10 millisecond latency but are more expensive than the machine vision camera approach we present here. That all said, these occurrences are fairly rare and default settings and parameters can be used when initially setting up the system.

**Table 2.** Latencies in the automated target chasing system.

|  |  |
|---|---|
| Latency for registered captures | 31 to 35 ms |
| Bonsai-Capture TTL latency | 28.96 to 29.27 ms (phase-lock R 0.992 to 0.999) |
| LED stamp jitter | 1.2 ms std; |
| Door reply, behind its TTL edge | 31 to 50 m. |
| Command to door pulse | Open: 7.24 to 7.48 ms; Close: 5.08 to 5.36 ms |
| Bpod pulse widths | 1000.08 ms ±6 µs; line 4: 43.83 ms ±16 µs |

This rig permits an examination of the behavioral and neural dynamics underlying target chasing and interception and can be expanded to multiple domains. What we present is the base condition of interception of randomly moving targets, but it should be noted that the current code and protocol can be modified for many use cases. Pursuit is a highly dynamic and ethologically-grounded behavior that involves continuous sensorimotor processing, planning, spatial awareness, value estimation, goal directed behavior, and decision making.

## ACKNOWLEDGEMENTS

We would like to thank Jack Carpino for developing the initial automated door design and Nethra Iyer for improvements to the motor system and lighting. This research was supported by NIH NINDS R00NS119665, DOE-ASCR CRCNS 0000285237, and the Whitehall Foundation. X.L. was supported by the Daryl and Marguerite Errett Discovery Award in Biomedical Research and a University of California President’s Postdoctoral Fellowship.

